# Kilohertz frame-rate two-photon tomography

**DOI:** 10.1101/357269

**Authors:** Abbas Kazemipour, Ondrej Novak, Daniel Flickinger, Jonathan S. Marvin, Jonathan King, Philip Borden, Shaul Druckmann, Karel Svoboda, Loren L. Looger, Kaspar Podgorski

## Abstract

Point-scanning two-photon microscopy enables high-resolution imaging within scattering specimens such as the mammalian brain, but sequential acquisition of voxels fundamentally limits imaging speed. We developed a two-photon imaging technique that scans lines of excitation across a focal plane at multiple angles and uses prior information to recover high-resolution images at over 1.4 billion voxels per second. Using a structural image as a prior for recording neural activity, we imaged visually-evoked and spontaneous glutamate release across hundreds of dendritic spines in mice at depths over 250 µm and frame-rates over 1 kHz. Dendritic glutamate transients in anaesthetized mice are synchronized within spatially-contiguous domains spanning tens of microns at frequencies ranging from 1-100 Hz. We demonstrate high-speed recording of acetylcholine and calcium sensors, 3D single-particle tracking, and imaging in densely-labeled cortex. Our method surpasses limits on the speed of raster-scanned imaging imposed by fluorescence lifetime.

## Introduction

The study of brain function relies on measurement tools that achieve high spatial resolution over large volumes at high rates^1^. Fluorescent sensors enable imaging of activity in large numbers of individual neurons or synapses on scales ranging from micrometers to millimeters^2–7^. However, the intact mammalian brain is opaque, and obtaining high-resolution images deeper than approximately 50 µm below its surface requires specialized techniques that are insensitive to light absorption and scattering. Two-photon imaging uses nonlinear absorption to confine fluorescence excitation to the high-intensity focus of a laser, reducing excitation by scattered light. Since fluorescence is generated only at the focus, scattered emission light can be collected without forming an optical image. Instead, images are computationally assembled by scanning the focus in space. However, this serial approach creates a tradeoff between achievable frame-rates and pixels per frame. Common fluorophores have fluorescence lifetimes of approximately 3 ns, requiring approximately 10 ns between consecutive measurements to avoid crosstalk^8,9^. The maximum achievable frame-rate for a 1-megapixel raster-scanned fluorescence image is therefore approximately 100 Hz, and is further limited in practice by factors such as photodamage and scanner bandwidth^1^. Biophysical events underlying neuronal communication, such as action potentials and neurotransmitter release^3,10,11^, occur on millisecond timescales. Kilohertz megapixel *in vivo* imaging would enable monitoring of these signals at speeds commensurate with signaling in the brain.

The pixel rate bottleneck can be avoided by efficient sampling^1,4^. When recording activity, raster images are usually reduced to a lower-dimensional space after acquisition, *e.g.* by selecting regions of interest^12,13^. Random-Access Microscopy^14–16^ samples that space more directly by only scanning targeted sets of pixels, with an access time cost to move between targets. When targets are sufficiently sparse, time saved by not sampling intervening areas significantly outweighs access times. Another efficient sampling approach is to mix multiple pixels into each measurement. Multifocal Multiphoton Microscopy scans an array of foci through the sample, generating an image that is the sum of those produced by each focus^17^. Elongated foci such as scanned Bessel beams^18–21^ collapse one dimension of the sample, generating projections of a volume at the rate of two-dimensional images. We refer to these and related approaches^4,22^ as “Projection Microscopy” because they deliberately project multiple resolution elements into each measurement. Projection Microscopy samples space densely, providing benefits over random-access microscopy in specimens that move unpredictably (e.g. awake animals or moving particles) or when targets are difficult to select rapidly (e.g. dendritic spines). However, recovering underlying sources from projected measurements requires computational unmixing^17^.

Computational unmixing is used in many imaging modalities to remove blurring due to finite resolution^23^, or to combine images having distinct optical transfer functions^24–27^. When imaging activity, pixels often contain signals from multiple sources, such as a neuron and surrounding neuropil, which analyses often model explicitly^12^. Unmixing is usually posed as an optimization problem, using regularization to impose known priors on the structure of recovered sources, such as independence^28^ or sparsity12. If the sources and measurements are appropriately structured, many sources can be recovered from relatively few mixed measurements^29^ in a framework called compressive sensing^30–33^. Highly coherent measurements (i.e. that always mix sources together the same way) make recovery ambiguous, while incoherent measurements can guarantee accurate recovery^29,34^. Tomography records linear projections of a sample along multiple angles. Tomographic measurements have low coherence because lines of different angles never overlap at more than one point. In medical imaging, compressive sensing using several tomographic angles has increased imaging speed and reduced radiation dose^33^.

Spatiotemporal patterns of brain activity encode sensory information, memories, decisions, and behaviors^35–37^. Neuronal activity exhibits transient synchronization at frequencies ranging from 0.01 to 500 Hz, giving rise to macroscale phenomena associated with distinct cognitive states^38^. In cortex, synchronous activity reflects attention^35,39^, task performance^39,40^, neuromodulation^41,42^, and gating of communication between brain regions^36,43^. Synchronous fluctuations in synaptic input are prominent during specific phases of sleep and anaesthesia^41^, but also present in awake animals^36,44^. Such patterns have been explored at the regional scale^45–47^, but activity patterns sampled by dendrites of individual neurons have not been recorded with high spatiotemporal resolution.

We developed a microscope that records high-resolution images spanning hundreds of micrometers at kHz frame-rates deep in the intact brain. This microscope combines benefits of random-access imaging and projection microscopy by performing tomographic measurements from targeted sample regions, enabling accurate unmixing when combined with priors. As priors for recovering neural activity, we use a structural image of the sample and a model of the fluorescent indicator dynamics, without regularization for sparsity. We used this microscope to record visually-evoked and spontaneous patterns of glutamate release on dendrites of isolated pyramidal neurons in mouse visual cortex, as well as for particle tracking, 3D imaging, dense samples, and other neural activity indicators.

## Results

### Scanned Line Angular Projection Microscopy

We built a microscope that scans line foci across the sample plane at four different angles (Fig. 1). Diffraction-limited line foci are simple to produce optically (Fig. S1), efficiently sample a compact area when scanned, produce two-photon excitation more efficiently than non-contiguous foci of the same volume, and perform low-coherence measurements. The resulting measurements are angular projections analogous to Computed Tomography^33^.

**Figure 1.**
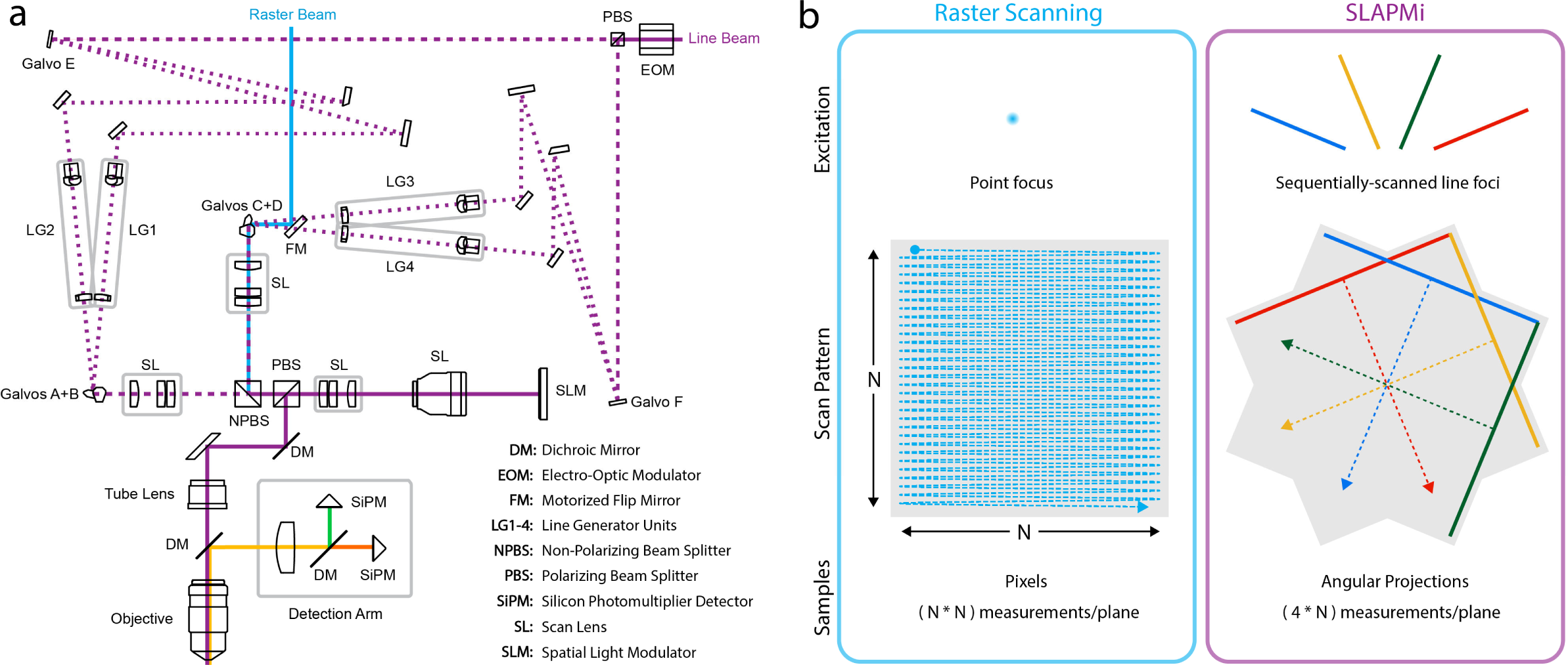
Scanned Line Angular Projection Microscopy. a)Optical schematic. SLAPMi directs a laser beam through four distinct excitation paths in sequence during each frame. Rapid switching between paths is achieved by an electro-optic modulator and polarizing beam splitter. Galvanometer mirrors (E,F) switch each beam again, producing four paths. Each path contains optics that redistribute the beam intensity into a uniform line of a different angle. Paths are recombined and scanned using two 2D galvo pairs (galvos A+B and C+D), and combined again at a nonpolarizing beamsplitter. The fully recombined beam is relayed and forms an image on a reflective SLM, which selects pixels in the sample to be illuminated, discarding excess light. SLM-modulated light returns through the relay and is imaged into the sample. A separate beam is used for raster scans. b) Raster scanning an (N-by-N)-pixel image requires N^2^ measurements. SLAPMi scans line foci to produce 4 tomographic views of the sample plane, requiring 4*N measurements.

We named this approach Scanned Line Angular Projection Microscopy (SLAPMi). SLAPMi can sample the entire field of view (FOV) with four line scans. The frame time is proportional to the diameter of the field of view in pixels *d*, in contrast to *d^2^* for a raster scan, resulting in greatly increased frame rates. For example, SLAPMi can image a 250×250 µm FOV with 200 nm resolution elements at a frame-rate of 1016 Hz, corresponding to over 1.4 billion pixels per second.

To reduce optical power at the sample and control the number of pixels mixed into each measurement, we included a spatial light modulator (SLM) in an amplitude-modulation geometry. This configuration selects a user-defined pattern within the FOV for imaging and discards remaining excitation light, making SLAPMi a random-access microscope (Movie 1). This avoids illumination of unlabeled regions, reducing excitation power in sparse samples, and allows users to artificially introduce sparsity into densely-labeled samples. Unlike other random-access methods, SLAPMi’s frame-rate is independent of the number of pixels imaged, up to the entire FOV.

### Particle Localization and Tracking

Localization of isolated fluorescent particles and tracking their diffusion or transport are used to monitor organelle dynamics, detect interactions between biomolecules, measure forces, and perform super-resolution imaging^48^. We demonstrated direct SLAPMi imaging of sparse samples by localizing and tracking fluorescent particles. Each particle produces a bump of signal on each scan axis corresponding to its position, and each frame consists of the superposition of signals for all particles in view (Fig. 2a,b). Using the microscope’s empirical measurement function, the maximum likelihood image can be recovered using Richardson-Lucy deconvolution^23,49^ (Fig. 2c,d). For large numbers of particles, maximum likelihood reconstruction produces spurious peaks in recovered images, which can be reduced by regularizing for sparsity (Fig. 2e).

**Figure 2.**
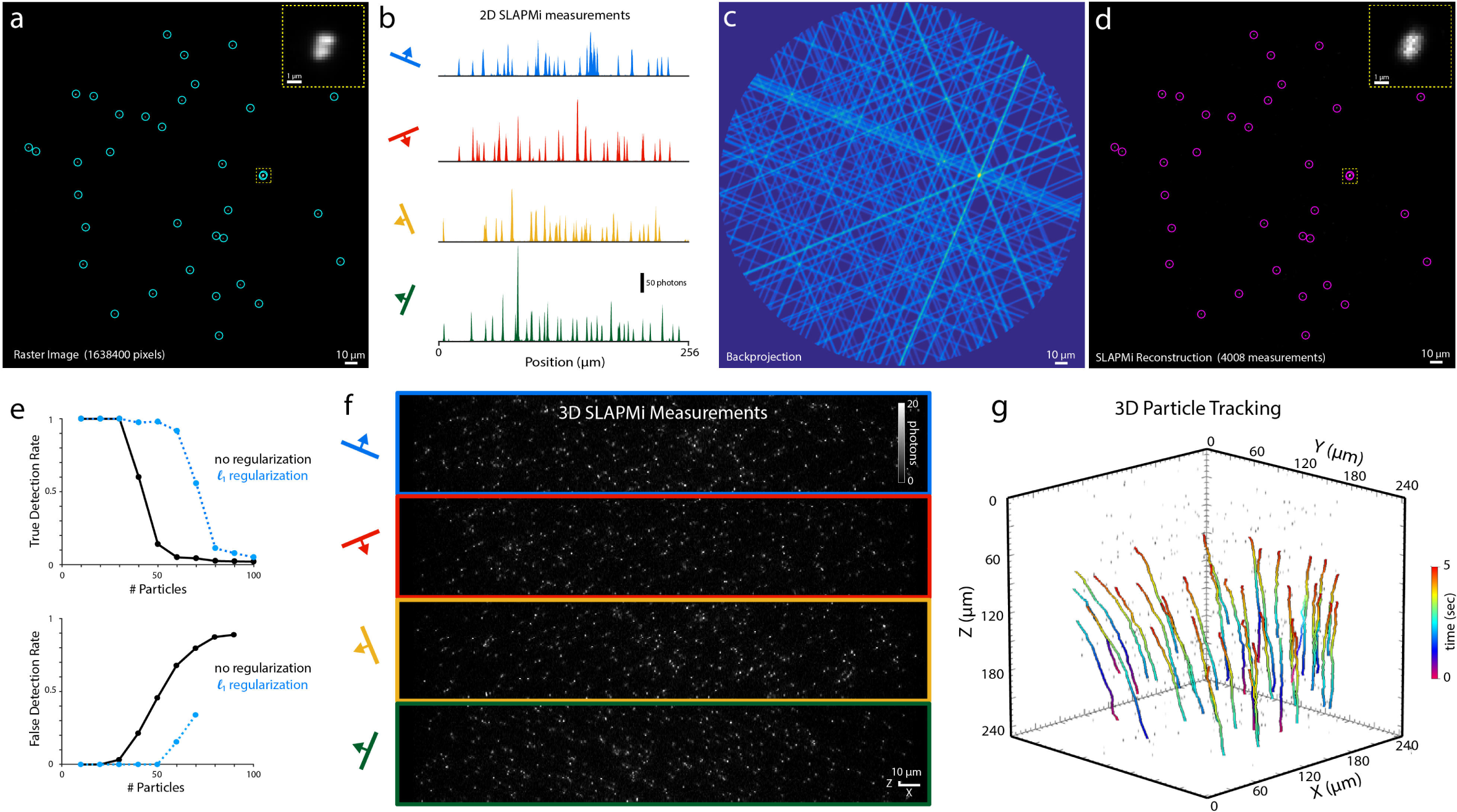
Particle tracking with SLAPMi. a) 2D raster image of fluorescent beads on glass. Circles denote detected particles (local maxima with intensity >20% of the global maximum). Inset shows zoom of a cluster of beads. b) 2D SLAPMi measurement of the same sample, consisting of projections along the four scan axes. c) Back-projection of the measurements in b. d) Recovered SLAPMi image with detections circled, demonstrating 100% detection with no false positives. e) Simulated detection performance for fields of view containing various numbers of point sources, with and without l_1_ regularization. (top) detection rate at 5% false positives; (bottom) false positives at 95% detection rate. f) 3D SLAPMi measurement of a 250×250×250 µm volume (90 planes) obtained in 89 ms. g) Paths of flowing particles in 3D SLAPMi video, tracked using ARIVIS software (Movie 3).

We performed 3D SLAPMi recordings of thousands of 500 nm fluorescent beads flowing in a 250 µm (diameter) x 250 µm (depth) volume (87 planes; 1016 Hz frame rate, 5080 measurements per plane, 10 Hz volume rate) (Fig. 2f, Movies 2,3). Particles were readily tracked in the resulting volume movies with existing commercial software (Fig. 2g).

### Computational Recovery of Neural Activity

To recover neural activity with SLAPMi, we adopted a sample representation in which spatial components (e.g. segments of dendrites) each vary in brightness over time^12,13^. The spatial components are obtained from a raster-scanned volume image, acquired in a manner that removes warping and motion artifacts. We segment this reference volume using a manually-trained pixel classifier (Ilastik^50^) and a skeletonization-based algorithm that groups voxels into contiguous segments (Fig. 3a, S4), resulting in up to 1000 segments per plane. Source recovery consists of determining the intensities **X** of each segment at each frame according to the following model:

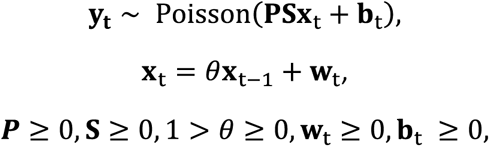

**Y=[y_1_,y_2_,…,y_T_]** are the measurements, each is a frame, **P** is the microscope’s measurement (projection) matrix, **S** encodes the segmented reference image, each **X**_i_ is the segment intensities on frame i, **b** is a rank-1 baseline, and θ implements the indicator dynamics.

**Figure 3.**
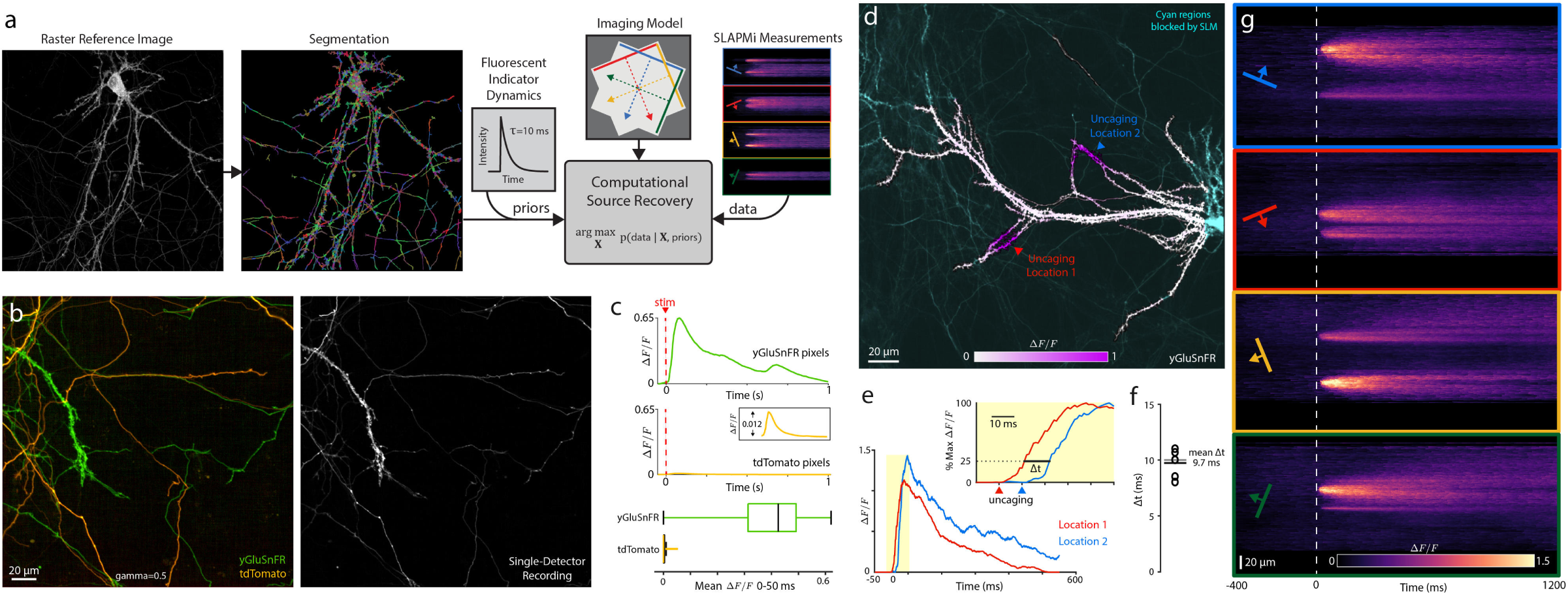
Validating SLAPMi activity imaging in vitro. a) Schematic of activity reconstruction. A segmented structural image and model of indicator dynamics are used as priors to constrain image recovery from tomographic measurements. b) Raster image of cultured neurons expressing either yGluSnFR or tdTomato. Only yGluSnFR responds to stimulation. (right) To test activity reconstruction, SLAPMi imaging was performed with one detector channel. c) (top,middle) Mean responses assigned to yGluSnFR- and tdTomato- expressing pixels. (bottom) Boxplot (quartiles) for distribution of mean ΔF/F_0_ 0-50 ms following stimulus for yGluSnFR- and tdTomato- expressing pixels. d) Uncaging glutamate at 2 locations on a yGluSnFR-expressing neuron. Saturation denotes maximum ΔF/F_0_ following uncaging. e) Responses at the two uncaging locations. Inset: Responses normalized to peak, demonstrating delay between responses. Arrowheads denote uncaging times at each location. f) Measured delays for 5 consecutive trials. g) SLAPMi measurements showing diffusion of signal following uncaging. Experiments shown are single trials without averaging.

This model imposes a strong prior on the space of possible recovered signals. It enforces that the only changes in the sample are fluctuations in brightness of the segments, corresponding to positive spikes and exponential decay dynamics. Motion registration is performed on **Y** and **S** before solving. The term that must be estimated is the spikes, **W =[w_1_,w_2_,…,w_T_]**. We estimate **W** by maximizing the likelihood of **Y** using a multiplicative update algorithm related to Richardson-Lucy deconvolution^51^. Importantly, in this paradigm, regularization is unnecessary because **S** has low rank by design. The solution is nearly always uniquely determined by the measurements, but adversarial arrangements of segments are possible.

We simulated SLAPMi imaging and source recovery while varying parameters such as sample brightness and number of sources, and assessed sensitivity to systematic modeling errors (Fig. S3). Simulated recovery was highly accurate at signal-to-noise ratios that match our experiments, and robust to errors in model kinetics, segment boundaries, or unexpected sources, but sensitive to motion registration errors. We assessed whether mixed measurements would introduce spurious correlations into recovered signals. Source-to-source correlations were recovered without bias (Fig. S3.7).

### Validation in Cultured Neurons

We performed SLAPMi imaging in rat hippocampal cultures, which facilitate precise optical and electrical stimulation in space and time. First, we co-cultured cells expressing the cytosolic fluorophore tdTomato with cells expressing the glutamate sensor SF-Venus-iGluSnFR^52^ (yGluSnFR; Fig. 3b). We imaged these cultures using one detector channel in which both fluorophores are bright, while electrically stimulating neurons to trigger yGluSnFR transients. TdTomato does not respond to stimulation, and any transients assigned to tdTomato-expressing cells must be spurious. This experiment tests the solver’s ability to accurately assign fluorescence in space, which we quantified using a separate two-channel raster image. Mean fluorescence transients assigned to yGluSnFR-expressing pixels were more than 52 times greater than mean transients for tdTomato (mean tdTomato 0.0078, maximum tdTomato: 0.0125, mean yGluSnFR 0.4192, maximum yGluSnFR 0.6525; Fig. 3c), with no tdTomato pixels having transients larger than 3% of the mean yGluSnFR amplitude, indicating that both frequency and amplitude of misassigned transients is low even in the densely intertwined cultures we imaged.

Second, we imaged yGluSnFR-expressing neurons while sequentially uncaging glutamate for 10 ms at each of two locations, to assess timing precision of recovered signals (Fig. 3d-g, Movie 4). Even though uncaging produced overlapping transients lasting hundreds of milliseconds, SLAPMi recordings reliably reported the 10 ms delay between these events (estimated delay 9.7 +/− 0.5 ms, N=5 recordings; Fig. 3f). Recordings showed the lateral diffusion of glutamate signals following uncaging^52^ (Fig. 3g).

### Glutamate Imaging *In Vivo*

We next recorded activity in dendrites and spines of pyramidal neurons in mouse visual cortex. Spines are specialized dendritic protrusions that receive excitatory glutamatergic input^6,53,54^. Pyramidal neurons are densely decorated with spines, which are often separated from each other and their parent dendrites by less than 1 µm^53^ (Fig. 4b), making spine imaging particularly sensitive to resolution and sample movement. SLAPMi’s high resolution and frame-rate allow precise post-hoc motion registration ideal for spine imaging (Fig. S5,S6).

**Figure 4.**
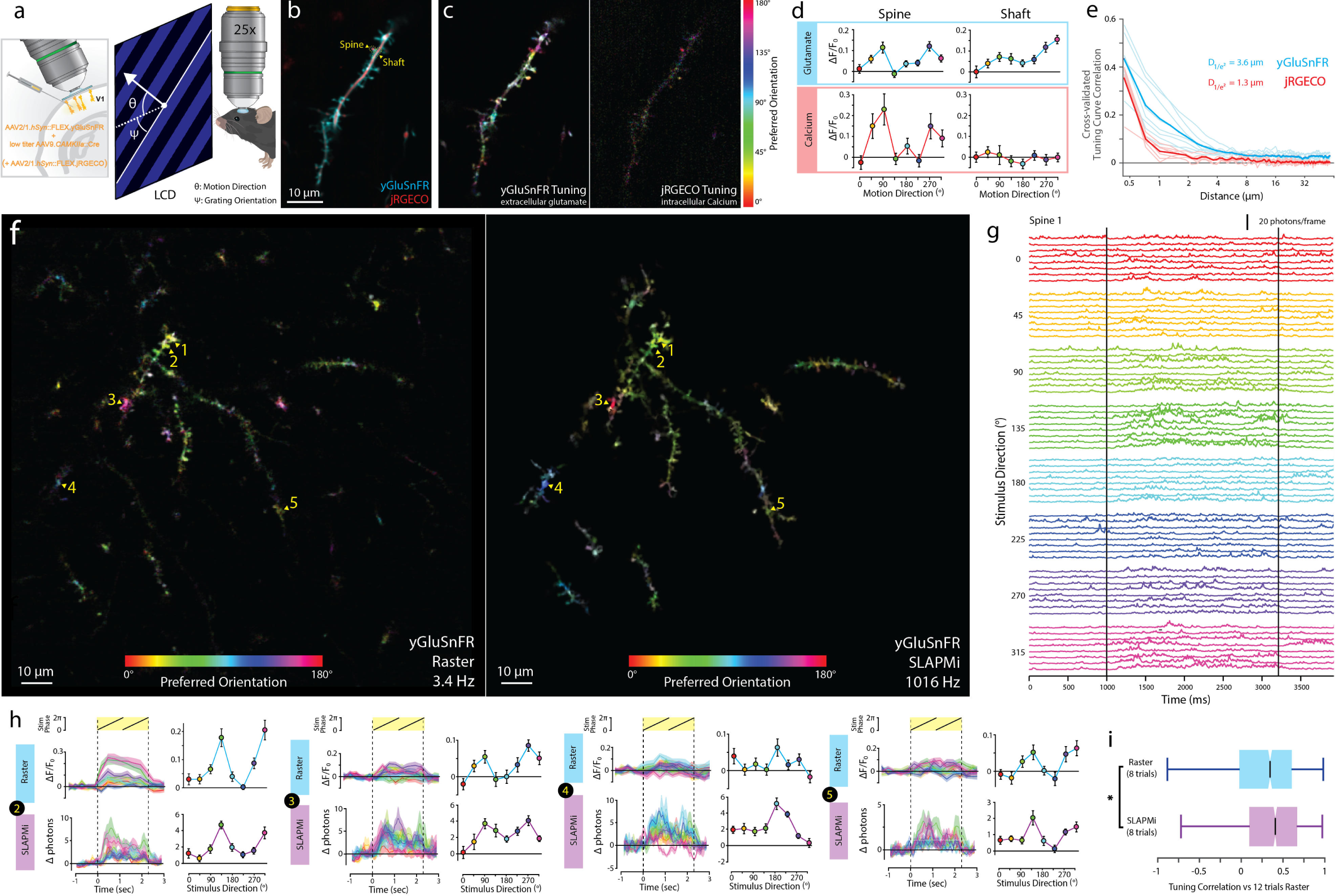
High-speed imaging of visually-evoked glutamate activity in cortical dendrites. a) Experimental design. Viruses encoding Cre-dependent yGluSnFR or yGluSnFR and jRGECO1a (b-e) mixed with low titer Cre were injected into visual cortex to sparsely label pyramidal neurons. We imaged responses to 8 directions of drifting grating motion stimuli. b) Raster image of dendrite in visual cortex co-expressing jRGECO1a (red) and SF-Venus.iGluSnFR.A184S (cyan). c) Pixelwise orientation tuning maps for each sensor. Hue denotes preferred orientation. Saturation denotes OSI. Intensity denotes response amplitude. d) Tuning curves for spine and adjacent dendritic shaft outlined in (b). N=20 trials for each direction. Errorbars denote S.E.M. e) Cross-validated correlation of pixel tuning curves, binned by distance, for simultaneously-recorded jRGECO and yGluSnFR signals. N=7 sessions, 3 mice, 20-25 repetitions of each stimulus/session. Dark lines denote means, light lines denote sessions. f) Pixelwise tuning maps for raster imaging (left; 20 repetitions per stimulus) and SLAPMi (right; 8 repetitions per stimulus) in yGluSnFR-expressing dendrites. g) SLAPMi recording (1016Hz) for a single spine over all trials grouped by stimulus type. h) Tuning curves and mean temporal responses for 4 spines recorded with raster imaging (20 trials per stimulus) and SLAPMi (8 trials per stimulus). Errorbars/shading denote S.E.M. i) Boxplots (quartiles) for distribution of correlations between tuning curves measured from 8 trials per stimulus of raster or SLAPMi and tuning curves from 12 raster trials per stimulus. *p<0.05, t-test.

Calcium transients within spines report NMDA receptor activation^54^ at synapses. However, NMDA receptors are activated by concurrent synaptic transmission and postsynaptic depolarization, not necessarily by synaptic transmission alone^55^. Moreover, spine calcium transients are triggered by other events, such as backpropagating action potentials, that may correlate with synaptic activity. Nonlinearities in calcium currents and fluorescent sensors^7^, and their slow timecourse, further complicate measurement of synaptic inputs using solely calcium transients. iGluSnFR enables detection of glutamate transients at individual spines in cortex^52^, potentially providing a complementary method to record synaptic activity.

To investigate the relationship between spine glutamate and calcium transients, we imaged dendrites of pyramidal neurons co-expressing yGluSnFR.A184S and the calcium indicator jRGECO1a^56^, while presenting lightly-anae sthetized (0.75-1% v/v isoflurane) mice with visual motion stimuli in eight directions. Vertebrate visual systems show orientation tuning, with response amplitudes often similar for a particular motion direction and its opposite^57^. Raster imaging indicated overlapping but different spatial response patterns between the two indicators (Fig. 4b,c). With trials containing dendrite-wide calcium transients removed, calcium responses were localized to a subset of dendritic spines, which showed large orientation selectivity indices (OSIs). At pixels with strongly-tuned calcium responses, glutamate and calcium tuning curves were highly correlated (Fig. S7). However, strongly-tuned glutamate responses were also present at spines with no calcium responses, and on dendritic shafts where no spine was visible (Fig. 4c,d). The spatial scale of glutamate responses was larger than that of calcium (1/e^2^ decay distance: 3.6 µm yGluSnFR.184S; 1.3 µm jRGECO1a, Fig 4e), indicating that yGluSnFR.184S is sensitive to the relatively low glutamate concentration changes outside the synaptic cleft. The glutamate affinity of yGluSnFR.A184S (7.5 µM^52^) is similar to that of the NMDA receptor (1.7 µM^58^). We therefore reasoned it could be used to investigate patterns of glutamate release relevant to NMDA receptor activation.

We interleaved SLAPMi (1016 Hz) and raster (3.41 Hz) recordings of FOVs containing hundreds of dendritic spines of isolated yGluSnFR.A184S-labeled layer 2/3 pyramidal neurons in visual cortex while presenting drifting grating stimuli. Both methods showed stimulus-evoked yGluSnFR transients on dendritic spines and shafts (Movies 5,6). Orientation tuning recorded with SLAPMi agreed well with much slower raster imaging (Fig. 4f,h), but SLAPMi revealed fluorescence transients too fast to be detected by raster imaging (Fig. 4g). SLAPMi was more accurate than raster scanning at predicting the tuning of dendritic segments as measured with larger numbers of raster trials (Fig 4i; SLAPMi r=0.35+/-0.02, Raster r=0.28+/-0.03; mean+/-SEM, p=0.015). To control for possible artefacts in imaging or source recovery, we generated a dead variant of yGluSnFR (yGluSnFR-Null) that lacks a periplasmic glutamate-binding protein domain. yGluSnFR-Null showed equivalent brightness and localization to yGluSnFR, but much less fluctuation, and no stimulus responses (N=4 sessions; Movies 6,7, Fig 5b).

Simultaneous recordings spanning large areas reveal how activity patterns travel over time^46^. In the absence of motion stimuli (Blank Screen), cross-correlations in yGluSnFR signals between pairs of dendritic segments were largest at short time delays, peaking at less than 10 ms regardless of distance over the 156 µm FOVs we imaged. In contrast, during motion stimuli, cross-correlations peaked at delays proportional to the distance between segments (linear fit 0.28+/-0.02 µm/ms, 95% c.i.; p<1e-11, N=10 sessions), indicating that activity patterns travel across cortical space as stimuli travel across the retina (Fig. 5c). This cortical magnification (12.7+/-0.7 µm/degree of visual field) was consistent with previous measurements over larger spatial scales^59^. Cross-correlations in yGluSnFR-Null recordings peaked weakly at a delay of 0 ms regardless of distance or stimuli.

**Figure 5.**
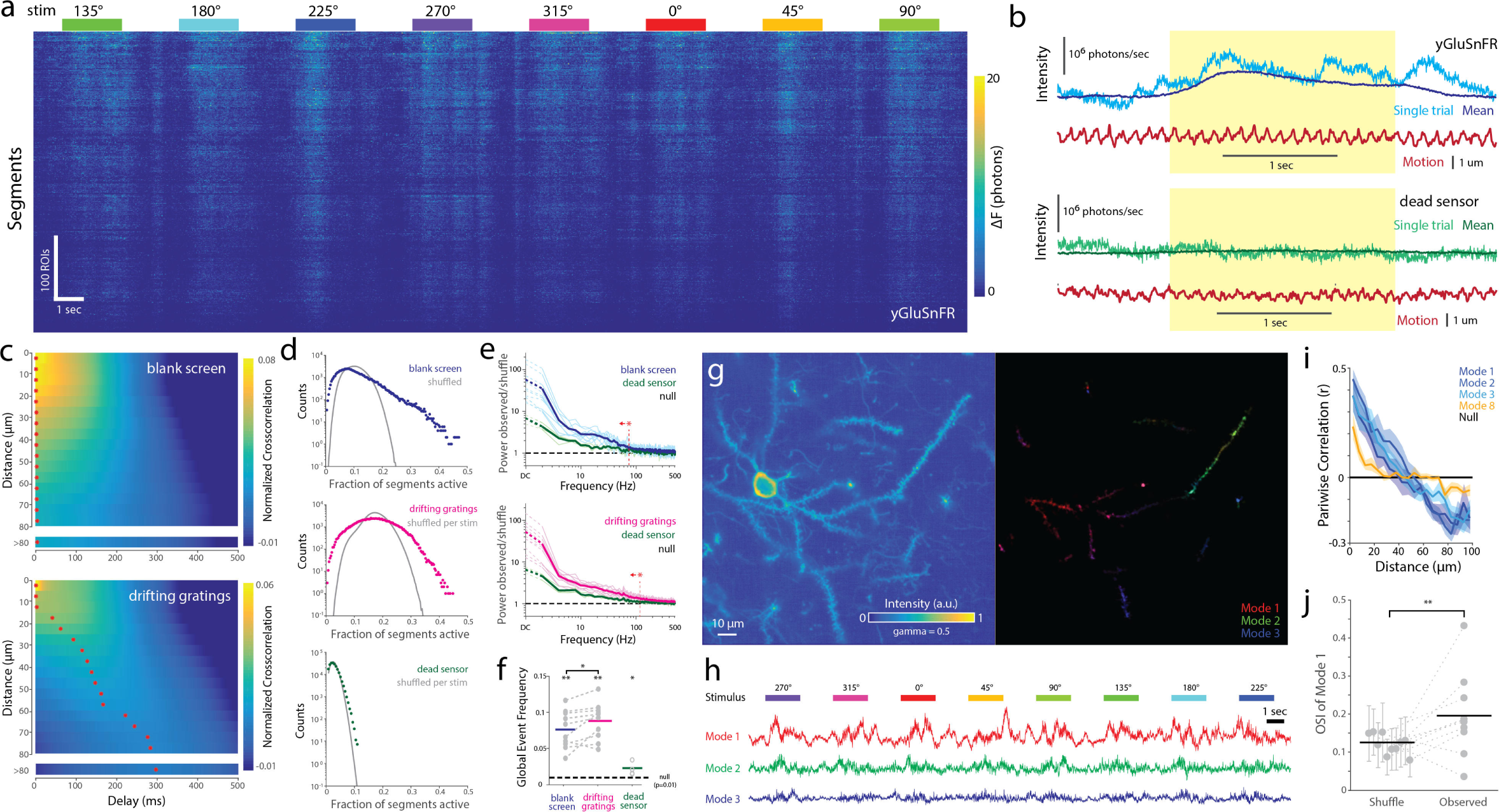
Spatiotemporal structure of glutamate transients in visual cortex. a) Recording of 460 dendritic segments exhibiting synchronized activity. b) Frame-integrated photon counts and estimated sample motion in yGluSnFR.A184S-labeled (top) and “dead” yGluSnFR-Null control (bottom). c) Mean cross-correlations between segment pairs, binned by distance, in the absence (top) and presence (bottom) of motion stimuli. N=10 sessions. Red stars denote cross-correlation maximum for each distance bin. d) Distribution of number of simultaneously active segments in the absence (top) or presence (center) of motion stimuli, and (bottom) corresponding distribution for dead sensor control. The null distributions are calculated by shuffling responses for each segment across trials of the same stimulus type. e) PSD of number of active segments, divided by corresponding PSD for the within-stimulus shuffled null, during blank screen and motion stimuli, for yGluSnFR and yGluSnFR-Null. Dark lines denote means, light lines denote individual sessions. Vertical red line denotes the minimum frequency bin at which yGluSnFR does not vary significantly from yGluSnFR-Null (t-test, p>0.05). f) Frequency of highly-synchronous activity (p<0.01 under null). *p<0.05, **p<0.01. N=10 sessions (yGluSnFR); 4 sessions (yGluSnFR-Null). g) Reference image (left), and spatial weights of the largest 3 NMF factors (right) for one session. h) Temporal components for the NMF factors in (g), over 8 consecutive stimulus presentations. i) Correlations between spatial component weights, binned by distance, for various NMF modes. Shaded region denotes standard error of the mean. N=10 sessions. j) Orientation tuning index of the largest NMF mode for 10 sessions, and null distribution obtained by shuffling stimulus labels. Errorbars denote standard deviation. **:p<0.01, Fisher’s combined test.

SLAPMi recordings revealed transients synchronized across dendritic segments on timescales faster than the 0.85 Hz stimulus frequency (Fig. 5a). This synchrony was visible in yGluSnFR recordings before source recovery, but absent in yGluSnFR-Null (Fig. 5b). To characterize synchronous activity, we selected dendritic segments with stimulus-evoked responses, and counted the proportion that were significantly active in each frame. The distribution of these counts can be compared to the null distribution obtained by shuffling activity traces for each segment among trials of the same stimulus type (Fig. 5d). Synchronization results in a higher probability of many segments all being active or all being inactive at the same time. yGluSnFR activity was significantly synchronized across segments both in the presence and absence of motion stimuli in all sessions recorded (N=10 sessions; Kolmogorov-Smirnov test, p<0.001). We compared the frequency spectrum of population fluctuations to the corresponding spectrum for the per-stimulus shuffled null. Activity was significantly more synchronized than yGluSnFR-Null controls at all frequencies below 117 Hz (during motion stimuli), or 73 Hz (blank screen) (t-tests, p<0.05; Fig. 5e). The frequency of highly synchronous events (p<0.01 under the null) was slightly increased in the presence versus absence of motion stimuli (8.8% vs. 7.6% of frames; N=10 sessions; paired t-test, p=0.03; Fig. 5f).

We investigated the spatial structure of synchronous activity by using principal component analysis (PCA) and nonnegative matrix factorization (NMF) to identify spatial modes (i.e. weighted groups of segments) that account for a large fraction of variance in our recordings. We used PCA to factorize recordings for each stimulus direction separately, after subtracting variance attributable to trial time, segment, or stimulus identities. The strongest resulting spatial modes were similar across all stimuli (Fig. S8), indicating that a small set of motifs accounts for large fluctuations in glutamate release onto cortical dendrites across stimulus contexts. The activity of these strongest modes was not significantly correlated to brain movement (Fig. S8). The largest PCA mode accounted for more variance in glutamate transients than did visual stimuli (10.3% vs 7.3%, adjusted R^2^; p=0.049, paired t-test, N=10 sessions). Using NMF to visualize spatial modes, we found that the strongest modes in each recording consisted of spatially-contiguous patches spanning 20-50 µm of dendrite (Fig. 5g,h). We quantified this spatial organization by calculating the correlation in mode weights between pairs of segments as a function of their distance. Across recordings, the strongest modes were spatially structured on a scale of approximately 50 µm, while weaker modes showed less spatial structure (Fig. 5i). Some spatial modes were strongly orientation-tuned while others were not; across sessions, the largest mode was significantly tuned (Fig. 5j; p=2.5e-4, Fisher’s combined test, N=10 sessions). From these results, we conclude that synchronous glutamatergic activity during anaesthesia is organized into patches within cortex, and that pyramidal neurons sample several such regions with their dendritic arbors.

### Other Imaging Modes

Recovering the voxel-space representation of a sample is not necessary in all experiments^32,60,61^. For example, NMF can be applied directly to motion-aligned SLAPMi measurements to extract responses and their corresponding projections on the four measurement axes (Fig. 6a-c). In another example, we used SLAPMi to identify punctate (^~^2 µm extent) locations of rapid (<30 ms) transients in cortex densely labeled with the acetylcholine sensor SF-Venus.iAChSnFR.V9 (yAChSnFR; Borden, Looger, et. al.; Manuscript in Preparation), 300 µm below the brain surface, triggered by electrical stimulation of the cholinergic nucleus basalis (Fig 6d-f). Transients could be identified on each projection axis, and backprojected to find their origin in pixel space, demonstrating that SLAPMi can record signals within densely-labeled samples without SLM masking if activity is sparse. These approaches retain the large sampling volume and optical resolution of SLAPMi without requiring a reference image, segmentation, or dynamics model.

**Figure 6.**
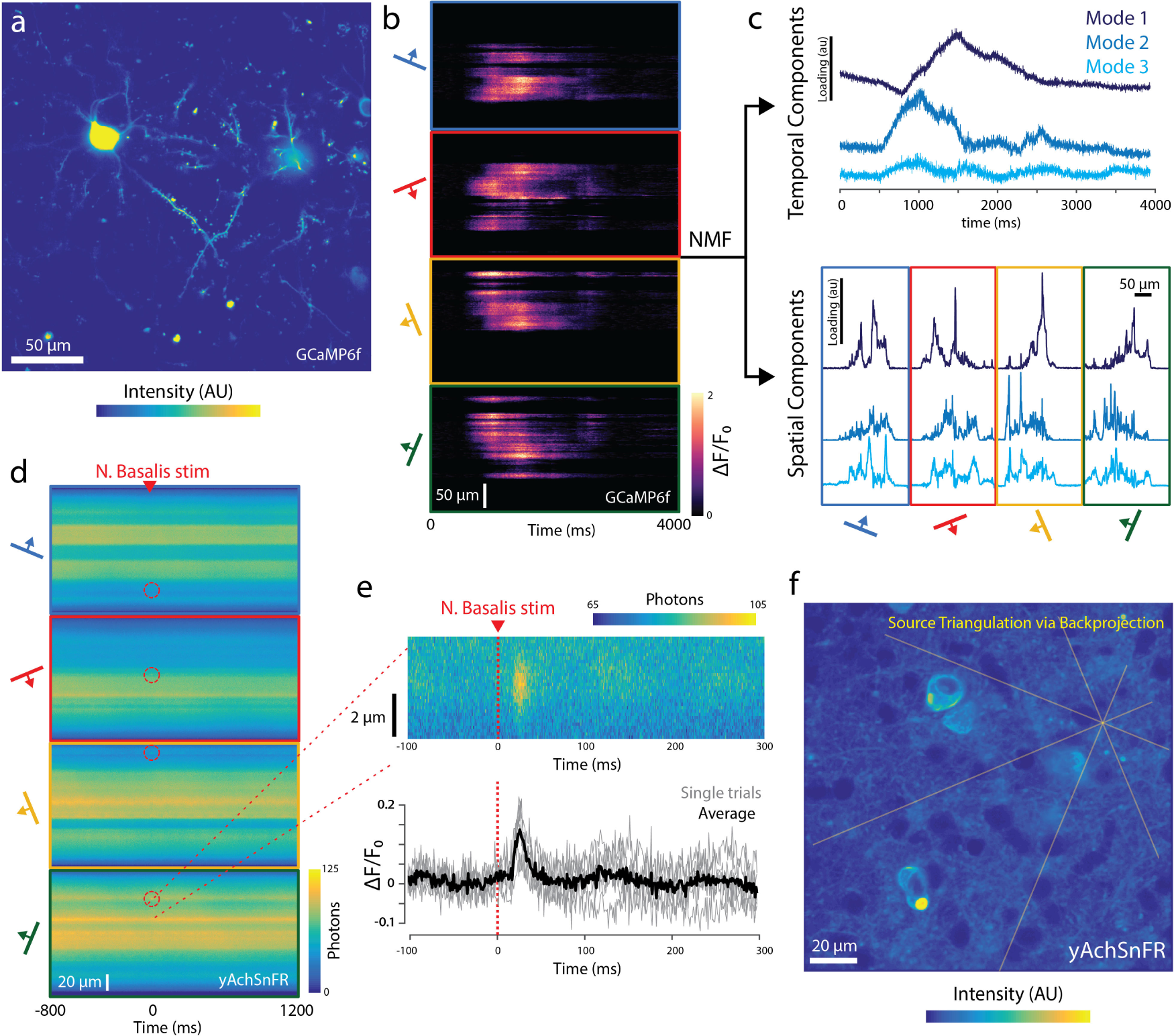
Alternative Imaging Modes. a) An imaging field containing multiple neurons expressing GCaMP6f. b) SLAPMi measurements from this field. c) NMF of the measurements reveals distinct spatial components (bottom), each with characteristic calcium indicator dynamics (top). d) SLAPMi measurements from an imaging field in visual cortex densely labeled with yAChSnFR, 300 µm below the brain surface. A single electrical pulse (500 µA, 1 ms) was delivered to nucleus basalis at time 0. Red circles highlight fluorescence transients visible in the four projection axes following stimulation. e) (top) Magnified view of stimulus-induced transient on one measurement axis. (bottom) ΔF/F0 traces for same region, over 6 consecutive trials. Dark line denotes mean, light lines denote trials. f) Backprojection of the locations in (d) onto raster image of the FOV reveals the spatial origin of the transients.

## Discussion

SLAPMi performs diffraction-limited *in vivo* imaging orders of magnitude faster than is achievable by point-scanning methods. This enables studies of rapid signals such as neurotransmitter release over FOVs spanning hundreds of micrometers. *In vivo* imaging techniques are becoming increasingly specialized, with different methods optimized for different experimental parameters^1^. Compared to acousto-optic deflector (AOD) random-access imaging, SLAPMi enables post-hoc motion compensation and higher frame-rates for large numbers of targets, but AOD-based imaging enables higher frame-rates for small numbers of targets. Compared to MMP and scanned Bessel beam approaches, SLAPMi benefits from lower coherence, higher frame-rates, and random-access excitation. SLAPMi’s mixed measurements limit the maximum number of sources to approximately 1000 per plane for the FOVs imaged here. As with any projection microscopy method, adversarial arrangements of sources over pixels can create ambiguities. In particular, regions where many distinct sources are packed closely together might be unrecoverable even when the total number of sources is small. Dim sources surrounded by very bright sources are recovered less precisely because shot noise associated with the bright sources dominates signals. SLM masking allows dense or bright regions to be blocked or dimmed, potentially ameliorating these issues. Compared to compressive sensing methods using sparsity priors, SLAPMi activity imaging avoids explicit regularization. Regularization parameter values can strongly impact reconstructed images, and can be difficult to select automatically. Spatial segmentation and dynamics are clearly-interpretable regularizers that take advantage of domain-specific knowledge.

Potential applications of SLAPMi include *in vivo* neurotransmitter imaging, large-scale volumetric calcium imaging, voltage imaging, recording rapid movements within muscles or vasculature, and tracking organelles or other particles in thick tissue.

## Concise Methods

### Optics

The microscope (Fig. 1) directs a single laser source (Yb:YAG, 1030 nm, 190 fs, tunable repetition rate 1-10 MHz; BlueCut, Menlo Systems, Germany) through four distinct excitation paths during each frame. An electro-optic modulator (EOM) rotates the beam polarization, enabling rapid switching between two paths at a polarizing beam splitter. A galvanometer mirror on each path again switches the beam between two paths. This configuration allows the beam to be sequentially directed onto the four paths with switching time limited by the EOM (<2 µs per switch). Only one path is illuminated at a time.

Each path contains a line generator assembly, consisting of an aperture followed by a series of cylindrical lenses, that redistributes the Gaussian input beam into a uniform line of tunable aspect ratio. The two pairs of paths are recombined at 2-dimensional galvo pairs. The resulting paths each pass through a scan lens and are combined using a nonpolarizing beamsplitter. This results in a loss of 50% of the beam but allows the transmitted beam to have a unique polarization. The beam is transmitted by a polarizing beam splitter and is relayed by scan lenses to form an intermediate image on the surface of a reflective liquid crystal spatial light modulator. Reflected light returns through the relay and the SLM-modulated pattern is directed to the sample at a polarizing beam splitter. The spatially modulated image is relayed by a tube lens and objective to the sample. Fluorescence emission is collected through the objective and reflected by a dichroic mirror to a non-imaging, non-descanned detection arm, as in conventional two-photon microscopes. We detect light with customized silicon photomultiplier modules (Hamamatsu). These detectors have high gain, high linearity, and exceptionally low multiplicative noise (Supplemental Methods, Fig. S9), making them ideal for parallelized imaging, where large numbers of detected photons arrive simultaneously. Wide-field epifluorescence illumination and camera detection (not shown) are coupled into the objective via a shortpass dichroic. For raster imaging, a different beam is introduced into the optical path by a flip mirror. The beam is scanned by one of the 2D galvo pairs as in conventional two-photon microscopes.

### Particle Tracking Experiments

Fluorescent beads (red fluorescent FluoSpheres, 0.5 µm; Molecular Probes, diluted 1:10000 in water) were sealed in a 2 mm diameter x 2 mm deep well capped by a coverslip. A 250×250×250 µm volume centered 150 µm below the coverslip was imaged using SLAPMi’s unidirectional Z-scanning mode at 10 Hz volume rate, 1016 Hz frame rate. Volumes were reconstructed with 30 iterations of 3D Richardson-Lucy deconvolution, using a projection matrix that accounts for the movement of the piezo. Tracking was performed with a linear assignment problem (LAP) tracking method in the commercial software ARIVIS.

### Hippocampal Culture Experiments

Rat hippocampal primary cultures were imaged 19 to 20 days after plating. For uncaging experiments, 3% of cells were nucleofected with a plasmid encoding CAG-SF.Venus-GluSnFR.A184V at time of plating. For experiments with tdTomato-labeled cells, a separate 3% of cells were nucleofected with a plasmid encoding CAG-tdTomato. In both cases, SLAPMi imaging was performed with a single channel using a 540/80nm filter. For uncaging experiments, 10 µM NBQX and 150 µM Rubi-Glutamate (Tocris) were added to the imaging buffer (145 mM NaCl, 2.5 mM KCl, 10 mM glucose, 10 mM HEPES, pH 7.4, 2 mM CaCl_2_, 1 mM MgCl_2_). Two-photon excitation power for culture experiments ranged from 39 to 42 mW at the sample. One-photon glutamate uncaging was performed with 420 nm fiber-coupled LEDs (Thorlabs M420F2). The tips of the fibers were imaged onto the sample plane through the same objective used for activity imaging. The two LEDs were activated at a current of 600 mA for 10 ms each in sequence.

### Surgical Procedures

All surgical procedures were in accordance with protocols approved by the HHMI Janelia Research Campus Institutional Animal Care and Use Committee (IACUC 17–155). We performed experiments with 19 C57Bl/6NCrl mice (females, 8-10 weeks at the time of the surgery). Each mouse was anaesthetized using isoflurane in oxygen (3-4% for induction, 1.5-2% for maintenance), placed on a 37°C heated pad, administered Buprenorphine HCl (0.1 mg/kg) and ketoprofen (5 mg/kg), and its head gently affixed by a toothbar. A flap of skin and underlying tissue covering the parietal bones and the interparietal bone was removed. The sutures of the frontal and parietal bones were covered with a thin layer of cyanoacrylate glue. A titanium headbar was glued over the left visual cortex. We carefully drilled a ~4.5 mm craniotomy (centered ~3.5 mm lateral, ~0.5 mm rostral of lambda) using a high-speed microdrill (Osada, EXL-M40) and gently removed the central piece of bone. The dura mater was left intact. Glass capillaries (Drummond Scientific, 3-000-203-G/X) were pulled and bevelled (30° angle, 20 µm outer diameter). Using a precision injector (Drummond Scientific, Nanoject III) we performed injections (30 nl each, 1 nl/s, 300 µm deep) into 6-8 positions within the left visual cortex. For sparse labeling experiments, the viral suspension was composed of AAV9.*CaMKII*.Cre.SV40 (Penn Vector Core, final used titer 1×10^8^ genome counts/ml) mixed with a virus encoding the reporter at 5×10^11^ gc/ml (one of: AAV2/1.*hSyn*.FLEX.GCaMP6f.WPRE.SV40 (Penn Vector Core); AAV2/1.*hSyn*.FLEX.iGluSnFR-SF-Venus.A184S (Janelia Vector Core), AAV2/1.*hSyn*.FLEX.iGluSnFR-SF-Venus.NULL (yGluSnFR-Null; Janelia Vector Core); AAV2/1.*hSyn*.FLEX.iGluSnFR-SF-Venus.A184V; AAV2/1.*hSyn*.FLEX.NES-jRGECO1a (GENIE Project vector 1670-115); AAV2/1.*hSyn*.FLEX.SF-Venus.iAChSnFR (Janelia Vector Core). For coexpression of jRGECO and yGluSnFR, we included both viruses at 5×10^11^ gc/ml each. The craniotomy was closed with a 4 mm round #1.5 cover glass that was fixed to the skull with cyanoacrylate glue. Animals were imaged 5-15 weeks after surgery.

### Visual Stimulation Experiments

A vertically-oriented screen (ASUS PA248Q LCD monitor, 1920×1200 pixels), was placed 17 cm from the right eye of the mouse, centered at 65 degrees of azimuth and −10 degrees of elevation. A high-extinction 500 nm shortpass filter (Wratten 47B-type) was affixed to the screen. Stimuli were drifting square-wave gratings (0.153 cycles per cm, 1 cycle per second, 22 degrees of visual field per second, 2 second duration) in 8 equally-spaced directions, spaced by periods of mean luminance, as measured using the microscope detection path. 0° denotes a horizontal grating moving upwards, 90° denotes a vertical grating moving rearwards. Mice were weakly anaesthetized with isoflurane (0.75-1% vol/vol) during recording. Anaesthetized mice were heated to maintain a body temperature of 37°C.

### Visualizations

All movies and recordings displayed are of single-trial recordings without averaging or additional filtering unless explicitly stated. See Supplemental Methods for details on data processing. Where noted, raster images have been square-root transformed (“gamma=0.5”) to better display dim features.

### Microscopy Software

Software was written in Matlab, C, and LabView FPGA. SLAPMi interfaces with ScanImage (Vidrio Technologies) to perform raster scanning. Plans for the microscope, instructions for use, example datasets, and code used in data acquisition, reduction, source recovery, analysis, and simulations are available from the authors (Update with Figshare DOI once available).

## Acknowledgements

This work is supported by the Howard Hughes Medical Institute. The authors thank Salvatore Dilisio, Na Ji, Aaron Kerlin, Justin Little, Christopher McRaven, Brett Mensh, Boaz Mohar, Manuel Mohr, Marius Pachitariu, Steven Sawtelle, Carsen Stringer, Srinivas Turaga and Ondrej Zelenka for contributions to this work.

The mouse image in Fig. 4a was obtained from the Database Center for Life Sciences (DBCLS TogoTV, © 2016; CC-BY-4.0 International license).

## Supplemental Methods

### Optical Methods

In the simplest scanning scheme, each frame consists of a single scan of each line orientation across the field of view (FOV). The maximum frame rate is determined by the cycle rate of the galvanometers, ~1300 Hz for a 250 µm FOV, limited by heat dissipation in the galvo servo controllers. We routinely image at 1016 Hz, to synchronize frames with the refresh rate of our SLM, which would otherwise produce a mild artifact. We also use tiled scanning patterns for efficiently scanning larger fields of view at only slightly reduced frame-rates (e.g. a 500 µm FOV at 800 Hz for tiling factor 2). The frame-rates achieved here could in principle be improved without substantial changes in the design, and are limited by a combination of scanner cycle rates and laser power. For an ideal scanner, the maximum frame-rate at any resolution would be achieved at a laser repetition rate of ~100 MHz (a 20x improvement over this work), above which the fluorescence lifetime would substantially mix consecutive measurements.

The number of paths was chosen to best suit tradeoffs between optomechanical complexity and acquisition speed (favoring fewer paths), versus benefits to source recovery (favoring more paths). For example, three measurement axes are necessary and sufficient to localize particles at low density, but additional axes are needed to resolve ambiguities at higher densities.

The line generator units (LG1-4) are used to impart a 1D angular range onto the incoming 2D Gaussian beam, such that when that beam is focused by a subsequent laser scanning microscope it makes a line focus with substantially uniform intensity distribution along the line, and diffraction-limited width in the focused dimension. The LG uses 3 cylindrical lenses. The first imparts a large, negative spherical aberration (in 1D) to the beam, which transforms the initial spatial intensity distribution into a uniform angular intensity distribution. The two subsequent cylindrical lenses refocus the light in 1D onto the 2D galvanometer mirrors. Thereafter, the microscope transforms the uniform 1D angular intensity distribution at the pupil into a uniform 1D spatial intensity distribution at the line focus at the sample. The relative lateral and axial translations of the first two cylindrical lenses allow an adjustment of the transform between the initial spatial intensity profile and subsequent angular intensity profile. Because spherical aberration alone cannot linearize an initial peaked power distribution over the entire initial distribution, a 1D mask is used at the beginning of the LG to remove the tails from the intensity distribution of the incoming beam. Because a sharp-edged mask at this location would create a 1D symmetric diffraction pattern, a mask with toothed edge profile was utilized. Alternately, the first two cylindrical lenses could be replaced by a suitable acylinder lens. Such a system would have fewer degrees of freedom to actively adjust the final line intensity distribution, but would allow an intensity transform function that can utilize the full tails of the initial spatial intensity distribution.

Axial scanning is produced by moving the objective with a piezo objective stage. SLAPMi is also compatible with remote focusing systems placed after the SLM in the optical path.

### Detectors

The parallel excitation used in SLAPMi can result in large numbers of emitted photons (from 0 up to ~400 in our experiments) arriving simultaneously in response to a single laser pulse. This necessitates a detector with large dynamic range, and higher photon rates favor detectors with low multiplicative noise. SLAPMi uses Silicon Photomultiplier (SiPM) detectors (MPPCs; Hamamatsu, custom part) for both raster scanning and fast line scan acquisitions. These detectors have extremely low multiplicative noise, sufficient gain to easily detect single photons, and are highly linear in their response within the range of photon rates we encounter, making the integrated photocurrent a precise measure of the number of incident photons (Fig. S9). The measured quantum efficiency of our detection path (for light originating from the objective pupil) is 32% at 525 nm and 20% at 625 nm, comparable to other microscopes at Janelia with the same detection geometry using photomultiplier tubes. Detector voltage is digitized at 250 MHz, synchronized to laser emission via a phase-locked loop. Photon counts are estimating by integrating photocurrent within a time window following each laser pulse and normalizing to the integrated current for a single photon.

### Actuator Control

To maximize the cycle rate of the galvanometers and piezo objective mount, we optimized the control signals used to command the manufacturer-supplied servo controllers for these devices. We optimized command waveforms iteratively, by measuring the error in the response waveform on each iteration, and using second-derivative regularized deconvolution to update the command signal in a direction that reduces that error, while maintaining smoothness. These calibrations take less than 30 seconds and are performed once per set of imaging parameters. With this approach, we were able to drive our actuators over 50% faster within our position error tolerance (<0.1% RMS error) than with methods that account only for actuator lag.

### Optimizations for Excitation Efficiency

Economy of illumination power is critical for biological imaging^62^, and conventional two-photon imaging is limited by brain heating under common configurations^63^. We were concerned that sample heating from light absorption could limit the practicality of two-photon projection microscopy methods such as SLAPMi, because higher degrees of parallelization require a linear proportional increase in power to maintain two-photon excitation efficiency. Higher degrees of parallelization also make source recovery more challenging by increasing background excitation and mixing between sources. In general, multiphoton projection methods benefit by using the lowest degree of parallelization compatible with an experiment’s required frame-rate^2,63^.

SLAPMi uses a customized Yb:YAG amplifier delivering high pulse energies (>9W @ 5MHz) at 1030 nm. The laser’s pulse compressor was tuned to achieve the minimum pulsewidth at the sample, ~190 fs at 5 MHz. We performed simulations to optimize the laser repetition rate given estimated nonlinear and known thermal^63^ damage thresholds of the sample. The optimal rate (data available on request) is approximately 5 MHz for single-plane imaging and 1 MHz for volume imaging in simulations we performed. To maximize excitation efficiency, each measurement consists of a single laser pulse.

For a given focal area within the sample plane, coherent line foci produce more efficient two-photon excitation than arrays of isolated points. This effect is partly because the edges of isolated points produce weak excitation. Line foci have a lower perimeter-to-area ratio than isolated points, making excitation more efficient.

The microscope contains a Spatial Light Modulator (ODPDM512-1030, Meadowlark Optics) that is used as an amplitude modulator to reject unnecessary excitation light. Cortical dendrites fill only a small fraction of their enclosing volume (<3% of voxels in our segmentations of single labeled neurons), allowing the majority of excitation light to be discarded. We retain a buffer of several micrometers surrounding all points of interest to guard against brain movement. The fraction of the SLM that is active depends on the amplitude of motion and the sample structure but is approximately 10% in most of our dendritic imaging experiments, substantially decreasing average excitation power. For sufficiently sparse samples, the SLM and other optimizations outlined above allow SLAPMi to use less laser power than full-field raster scanning (Fig. S2).

The combination of SLM masking and scanning used to vary the excitation pattern in SLAPMi produces much higher pattern rates (5 MHz) than can be achieved with SLMs alone (<1 kHz), and more control over those patterns than scanning alone. This approach could be used similarly with other projection microscopy approaches, and is easiest to implement if all excitation foci can be made to lie in the plane of the SLM.

### Reference Images

Accurate registration and source recovery require minimally warped reference raster images. The software that operates SLAPMi interfaces with ScanImage (Vidrio Technologies) to perform all point-scanning imaging. Raster images can be warped by many factors, including nonlinearity in the scan pattern, sample motion, and the ‘rolling shutter’ artifact of the raster scan. These errors must be corrected in the reference stacks to perform accurate source recovery. We estimate and compensate for warping by collecting two sets of reference images interleaved, one with each of the two galvos acting as the fast axis. We assume that motion is negligible during individual lines along the fast axis (~300 µs), and that the galvo actuators track the command accurately along the slow axis. These assumptions allow recovery of unwarped 2D images by a series of stripwise registrations. The axial position of the image planes is estimated by alignment to a consensus volume.

### Segmentation

The reference stack must be segmented into compartments such that the number of compartments in any imaging plane is less than the number of measurements. We use a manually trained pixel classifier (Ilastik^50^ in autocontext mode) to label each voxel in the volume as belonging to dendritic shaft, spine head, other location within a labeled neuron, or unlabeled background. We do not deconvolve the reference stack or attempt to label features finer than the optical resolution. Instead, we label features at the optical resolution and design the projection matrix **P** such that it accounts for the transformation from the point spread function of the raster scan to that of each line scan (Fig. S1). We use a skeletonization-based algorithm to agglomerate labeled voxels into short segments of roughly 1.4 µm, which approximately correspond to individual spines or short segments of dendritic shafts. Segments that are predicted to produce very few photon counts (after considering SLM masking) are merged with their neighbors. Source recovery performs well when there are less than 1000 segments per plane. Most fields of view contain 400 to 600 segments.

### Measurement Matrix

The microscope’s measurement (projection) matrix **P** is measured in an automated process using a thin (<<1 µm) fluorescent film (see `measure_PSF` in the software package). Images of the excitation focus in the film, collected by a camera, allow a correspondence to be made between the positions of galvanometer scanners and the location of the resulting line focus. The raster scanning focus is also mapped, allowing us to create a model of the line foci transformed into the space of the sample image obtained by the raster scan. The camera is unable to resolve the shape of the excitation focus of the line along the shortest axis (the R-axis). The point spread function on the R and Z axes is measured by scanning sparse fluorescent beads, but this measurement is not needed to generate **P** because we model SLAPMi measurements as a fixed convolution of the raster point spread function, which is already encoded in the raster reference image.

SLAPMi achieves the same lateral resolution as point scanning two-photon, but axial sectioning is slightly poorer in SLAPMi because light is focused on only one axis to produce each line (full-width-half-maximum 1.62 µm vs. 1.46 µm for point focus, Fig. S1). This increased depth of field can in some cases be of benefit as it reduces effects of axial motion.

### SLAPMi Data Reduction

Raw data recorded by SLAPMi consists of detector voltages and galvo position sensor voltages (digitized at 250 MHz, synchronized to the laser output clock via phase-locked-loop), and auxiliary analog inputs (such as photodiode voltages to monitor visual stimulus retrace times, and piezo objective stage position sensors for 3D experiments) digitized at up to 10 MHz. Data reduction converts these raw voltages into frames, containing intensities (in units of photons) assigned to a set of reference galvo positions, and other ancillary data. Recording the galvo positions and performing this registration compensates for drift in the galvo path during high-speed recordings that would otherwise contribute to noise. For details, see `SLAPMi_reduce` in the software package.

### SLAPMi Motion Registration and Alignment

Recorded SLAPMi data are spatially registered to compensate for sample motion. As with raster imaging, translations of a single resolution element can be sufficient to impact recovery of activity in fine structures, necessitating precise registration (Fig. S3.7). Frame-by-frame alignment (Fig. S5) is first performed by cross-correlation of each projection axis on each frame to a consensus centroid of the measurements. This corrects for rapid in-plane motion of the sample. The stabilized recording is then averaged over time in blocks, and each block is aligned for 3D shift relative to the reference image (Fig. S6). This registration is performed by identifying the 3D translation of the sample that maximizes the sum of correlations (or optionally, minimizes Dynamic Time Warping distances^64^) between the recorded signal and the expected measurements on each of the four scan axes. If the SLM is not used, this objective is well approximated using cross-correlations that can be rapidly computed. If the SLM is used and a significant amount of sample intensity crosses to borders of masked regions during motion, cross-correlations are not effective and we instead perform an iterative multiscale grid search using the full measurement matrix.

### Analyses of *in vitro* recordings

In Fig 3c, source recovery was performed on Channel 1 of the SLAPMi recording and projected into two dimensions using the axial mean projection of the segmentation matrix S. ΔF/F_0_ was fit for each voxel in this projection. Fig. 3c shows the mean ΔF/F_0_ for the period 0-50ms after the stimulus, for the greenest 1/3 of pixels and the reddest 1/3 of pixels in the reference image within the segmentation. In Figure 3e-f, we selected the dendritic segment closest to each uncaging location, which were registered relative to the scanner coordinates by imaging in a thin fluorescent sample. Δt was calculated as the difference between times at which each trace crossed 25% of its maximum within the 150 ms period following the first uncaging event. Figure 3f shows Δt values from single trials across 5 experiments.

### Analyses of *in vivo* recordings

Raster activity recordings were aligned with custom MATLAB code that compensates for bidirectional scanning artifacts, warping of galvo scan paths, and rolling shutter movement artifacts using a series of stripwise registrations, included in the published software package (`alignScanImageData` and related functions). For simultaneous recordings of jRGECO and yGluSnFR (Fig. 4), the two recording channels (540/80 and 650/90 bandpass filters) were linearly unmixed with the mixing coefficients 0.069 (yGluSnFR->jRGECO bleedthrough) and 0.058 (jRGECO->yGluSnFR bleedthrough), estimated using robust regression against pixel intensities in the aligned average images.

The method of calculating F_0_ was different for different imaging methods. In raster recordings, F0 for a region of interest was calculated for each stimulus presentation as the mean intensity in that region over 3 frames prior to the stimulus. For SLAPMi recordings, the terms **X** in the activity model can be directly interpreted as the ΔF/F_0_ values for each segment. The baseline fluorescence is accounted for by the time-varying baseline term **b**, which was fit in the space of the measurements based on the cumulative minimum of the regression of the observed against the expected measurements (see `SLAPMi_solve`). For plots of ΔF/F_0_ in the measurement space (Fig. 3g,6b), a time-varying F0 was fit based on the cumulative minimum of the raw intensity for each row of **Y**, and ΔF/F_0_ was computed, temporally smoothed using non-negative deconvolution^65^ and downsampled for display.

In Fig. 4e, correlations in pixelwise tuning curves were calculated by cross-validation; the dataset for each session was dividing into two halves and tuning curves were calculated for each pixel. For each pixel, the tuning curve from one half of the data was correlated against tuning curves obtained from the other half of the data for all other pixels. This was done for 5 different random splittings of the data, and results averaged, for each session. This approach avoids introducing correlations due to trial-to-trial variations.

Orientation selectivity indexes (OSIs) (Fig. 4b,5c,7d) were calculated from the 8-point tuning curves (*i.e.* the mean response during the stimulus period for the 8 stimuli, minus the mean response during 1 second prior to the stimulus) for SLAPMi and raster recordings in the same way. The 8 stimulus angles were mapped onto four equally-spaced angles (*i.e* 0,90,180,270,0 90,180,270 degrees), multiplied by response amplitude, and vector summed. The preferred orientation (hues in Fig. 4b,5c) is the angle of the sum vector. The OSI is the length of the sum vector, divided by the sum of the lengths of the input vectors. It has a value in [0 1] which is 1 when there is a response to only one of four orientations, and 0 when responses are uniform. The response amplitude is the 2-norm of the 8-point tuning curve.

Movies 6 and 7 are rendered as approximate Z-scores, computed as F/sqrt(F_0_+3/8), i.e. the measured photon rate (in units of photons/frame for each dendritic segment) divided by the square root of the baseline photon rate for each segment plus a small background (3/8). The samples in Movies 6 and 7 had similar labeling brightness and were imaged at the same laser powers prior to the SLM, produced similar mean photon rates per dendritic segment (14.5 photons/segment, Movie 6; 18.2 photons/segment, Movie 7), and were reconstructed using the same solver parameters.

In Figure 4i, Raster recordings were aligned to the SLAPMi reference image, and dendritic segments were projected into the plane of the raster recording to generate regions of interest for correlation analyses. Only segments with components within the plane of raster imaging were included in analysis. Tuning curves were obtained for 8 randomized trails, and compared to the remaining 12 trials of the raster recording. SLAPMi tuning curves (8 trials) were compared to the same 12 trials for each randomization, and the distribution of correlations for each method is plotted.

For SLAPMi population analyses (Fig. 5), we analyzed all segments with mean brightness greater than 1.5 photons per frame, for which no more than three trials of data were lost due to motion (“selected segments”). Motion censoring by SLAPMi is performed on the basis of goodness-of-fit of the SLAPMi 3D alignment method (`SLAPMi_PSXdata` and related functions). In the experiments shown here, only two trials across all datasets contained unacceptable sample motion and were censored. Additionally, for calculating stimulus tuning, we censored the first three stimulus presentations of SLAPMi recordings in each session.

In Fig. 5c cross-correlations in activity traces were calculated between all pairs among selected segments in each trial, normalized to the geometric mean of the variances for each pair, and averaged across trials and sessions for each distance bin weighted by the product of the two mean brightnesses. See the published code for more details. In Fig. 5df, we labeled each segment in each frame as active or inactive by high-pass filtering at 4 Hz and thresholding at 3.5 standard deviations of the bottom 80% of the data above the median. We analyzed the distribution of the number of active segments across frames. The null distribution was calculated by randomly shuffling each trial for each segment across the 8 trials of the same stimulus type for the same segment, and calculating the same distribution. This method controls for synchronization due to structure in the stimulus and shared stimulus tuning. In Fig. 5f, we calculated observed rates of events exceeding the p=0.01 level of the null distribution, and used Z-tests to compare to the known null rate (0.01). We used paired t-tests to compare the two stimulus conditions to each other.

In Fig. S8, spatial modes were calculated by PCA over concatenated trials of each stimulus after subtracting the mean response to that stimulus for each recorded segment. We report the correlations in the spatial weights of the modes across segments. Note that the precise ordering of modes can vary slightly across stimuli despite a one-to-one correspondence, such that *e.g.* the second largest mode of a given stimulus might correlate highly to the largest mode of all others. For visualization in Fig. 5g spatial modes were calculated by NMF, to overcome the orthogonality constraint imposed by PCA. Because NMF performs a strictly positive factorization, subtraction of mean responses was not performed, and modes were instead calculated only from fluctuations during the blank screen period. The spatial appearance of modes obtained by NMF was qualitatively similar to the positive components of modes obtained by PCA from the complete datasets after stimulus subtraction. The NMF modes were then used to obtain temporal components for all frames in the stimulus-subtracted data by regression (Fig. 5h). In Fig. 5i, we report the correlation in the spatial weights of pairs of segments, where the distance between the segments falls into the specified distance bin, for each mode. This correlation is positive for a given distance if segments spaced by that distance tend to have the same weight. It is zero if weights are independent and identically distributed (i.e. unstructured in space).

In Fig. 6b-c, SLAPMi measurements were aligned by crosscorrelation to a concensus mean on each measurement axis, and NMF was performed directly on the measured photon counts. In Fig. 6f, events were localized manually on the four measurement axes and the measurements corresponding to the selected locations were backprojected into the pixel space.

## Supplemental Notes

### Notations

Throughout the paper we have adopted the following notations:

Vectors and matrices are denoted by bold lowercase and capital letters respectively. Vectors are assumed to be column vectors.

For a vector we denote its transpose by V^2^. 1 and 0 are used to denote the all ones and all zeros vectors respectively. We denote the Hadamard (elementwise) product of two vectors (matrices) of the same size by ⊗ and ⊘ respectively. By 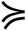 we denote elementwise inequality between two vectors. For a vector **V = [v,_1_,v_2_,…,v_n_]** we denote its 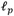 norm by: 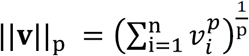. For a matrix **W = [w_1_,w_2_,…, w_T_]** we denote its 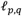 norm by 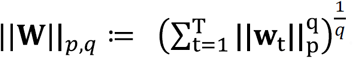.

### Solver

The solver performs a constrained maximum a posteriori (MAP) estimation on the innovations ( w_*t*’s_) in the state-space model given by:

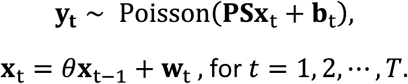

We set

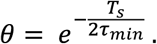

Where *T_s_* is the sample time and τ*min* is the fastest decay constant of the indicator being imaged. All terms are constrained to be nonnegative. We solve for **W** by minimizing the Poisson loss objective function:

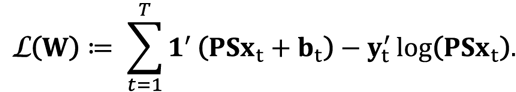

We have also implemented several priors, including penalization of *l_p,q_* norms of the innovations, which were not used in this study. We refer to (^51^) for details on how to incorporate regularization into this solver. Since that work, we have altered or improved several features, described below.

The solver performs a modified form of Richardson-Lucy deconvolution (RL) that incorporates the autoregressive (AR(1)) dynamics of the model (“dynamic RL”). RL iterations converge to the maximum likelihood solution to unmixing problems under Poisson sampling statistics. This is done by iteratively multiplying an estimator by the ratio of two positive terms, the difference of which is the gradient of the error. We adapted RL iterations by computing the gradient of the objective function in our dynamic model with respect to the innovations (**W**), splitting this into two positive terms, and forming multiplicative updates analogous to RL. Solving with respect to the nonnegative innovations enforces the desired dynamics. This approach is equivalent to augmentation of the measurement matrix (**PS**) with AR dynamics and solving the RL iterations for the innovations. Utilizing the time-invariance of **PS**, this can be done efficiently via RL updates followed by filtering operations using the AR dynamics. The partial derivative of the objective function with respect to **W**_t_ is:

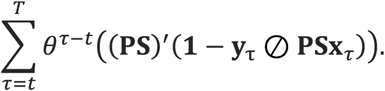

Therefore, the positive and negative terms of the update are:

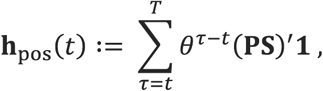

and

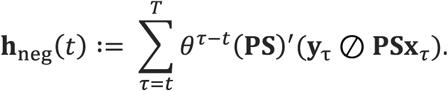

This constitutes the update rule for dynamic RL:

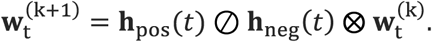

Additionally, we constrain the spikes not to change by more than a factor of 10 on each update, by clipping the gradients to satisfy 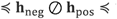. The solver is initialized with a positive constant **X**.

Updates following the direction of the gradient fail to fully account for the properties of the AR(1) model in a small number of iterations, namely the conditional independence of elements of **X**_t ′ > t_ on elements of **w_t_** given intervening elements of **X** (see ^65,66^). Because of this property, no **w_t_** need be made larger than the value that maximizes the likelihood at the t^th^ timepoint, because any further increase to **w_t_** could instead be applied to **X**_t ′ > t_ to better effect. The update that considers the gradient at the t^th^ timepoint alone is the unmodified RL iteration. We therefore use a modified update rule that is the minimum of the dynamic update and that of unmodified RL. This update rule alters the direction of update steps relative to dynamic RL but has the same stationary point.

In all datasets presented in this paper we estimated the baseline **b** from a minimum of smoothed measurements (**y**_t_′s), yielding a rank-one estimate. The baseline was assumed to be fixed and no updates were applied. We include options to use a fraction of this baseline or update it using RL type updates.

1. Maximum likelihood solutions are known to amplify noise, and RL reconstructions are typically terminated after relatively few (<100) iterations to compromise between degree of unmixing and unwanted artifacts^67^. The number of iterations performed similarly affects our solver. However, we have implemented two methods to suppress this effect: Optionally, we allow a regularization term to impose prior knowledge of the indicator to constrain the -norm of **X**. For example, an infinity norm can be used to constrain maximum 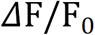 to a known maximum for the indicator. In particle localization, a small 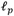-regularization (*p* ≤ 1) was helpful in pruning artifacts.
2. Optionally, we allow damped RL iterations^67^ which flatten the Poisson log-likelihood for smaller photon counts, reducing the size of update steps in these regions. For the activity recordings presented in the paper we used a damping parameter of 1.5, which reduces steps in regions where the difference between measurement and reconstruction are smaller than 1.5 σ.

### Inputs to the solver

The solver has few user-adjustable parameters, listed below:

1. The decay time constant: a physical property of the indicator, which needs to be specified. This time constant determines the AR parameters used by the Dynamic RL updates. A decay time constant of 0 corresponds to regular RL iterations.
2. Damping parameter: residuals less than the damping parameter standard deviations will have less effect on updates. Typically, the damping parameter is chosen in the range 0-2.
3. Minimum and maximum number of iterations of the algorithm. Typical values are in the range 50-200.
4. Dark noise level: in photons, added to all expected rates
5. Stimulus artifact: to include a stimulus artifact in the rate model used for the two-spot uncaging datasets, set this to the approximate artifact length. The artifact will be automatically detected.
6. Clip Gradients: if set to true it will clip multiplicative updates to the range [0.1, 10] per iteration.
7. Off-focus spikes: if set to true allows fitting of spikes for scattered excitation light, useful for deep *in vivo* imaging.
8. Baseline multiplier: fraction of the minimum of data used as the estimate of the baseline.
9. Reconstruction algorithm: Richardson-Lucy iterations or multiplicative nonnegative least squares.

Parameters 4,6,7,8, and 9 were not altered throughout activity imaging experiments and are provided primarily for future applications. Parameter 5 specifies an augmented model used in only one experiment. Parameter 1 should be set appropriately using a priori knowledge of the indicator, or slightly underestimated if uncertain. Default values of the other parameters are suitable for most datasets.

Summary of parameter values for datasets in this paper:

|  | Particle Localization<br>(no dynamics) | In-Vitro Red/Green | In-Vitro 2-spot<br>uncaging | In-Vivo iGluSnFR |
| --- | --- | --- | --- | --- |
| Decay time constant | N/A | 50 ms | 50 ms | 10 ms |
| damping | 0 | 1.5 | 1.5 | 1.5 |
| Minimum/maximum iterations | 100-100 | 50-100 | 50-100 | 50-100 |
| Dark noise | N/A | 0.01 | 0.01 | 0.01 |
| Stimulus artifact | N/A | N/A | 10 ms | N/A |
| Gradient clipping | false | true | true | true |
| Baseline multiplier | N/A | 1 | 1 | 1 |
| Off-focus spikes | N/A | false | false | True |
| Reconstruction algorithm | RL | Dynamic RL | Dynamic RL | Dynamic RL |

The solver requires a 2017a or later release of MATLAB to run properly. In order to run the solver you need to have all the functions in the /slapmi directory on your MATLAB path.

The main function for the solver is `SLAPMi_solve`, which upon running will prompt you for the problemdata directory (can choose multiple problemdata files) and the desired solver option as a graphical user interface which is set to default options suitable for most datasets.

The “.psf” files corresponding to the microscope calibration at the time of acquisition should either be on your path or in the same folder as the corresponding problemdata file(s). “.psf” files are generated by the microscope during alignment, by the function `measurePSF`.

Problem files are generated by the function `SLAPMi_PSXdata`, which prompts for input of a reference raster volume (generated by `SLAPMi_REFim`), an Ilastik project (.ilp) file, and reduced raw data from the microscope (generated by `SLAPMi_reduce`).

The solver will generate a child folder /Recon where the reconstruction files are saved as well as a /ReconMovies folder where the reconstructed videos are stored. (if `opts.plot = true`)

A typical instance of running the solver begins with the function call:

~~~
sys_recon = SLAPMi_solve(opts)
~~~

The results can be visualized by running

~~~
SLAPMi_plot(sys_recon)
~~~

Which will automatically generate and save the reconstruction movie in the reconMovies folder.

The output structure `sys_recon` contains the following 5 fields:

`sys_recon.opts`: which includes user input options.

`sys_recon.input`: which includes problemdata and scandata inputs to the solver

`sys_recon.output`: which includes the outputs of the solver including the estimated spikes and segment activities (`sys_recon.output.spikes` and `sys_recon.output.F` respectively).

`sys_recon.solver_params`: internal parameters which are used by the solver. These parameters are fixed and a function of the problemdata and do not require modification.

`sys_recon.control_params`: control parameters for checking the validity of estimates. These parameters include the objective function (used to determine convergence), regularizations (if used), information about the solution at each iteration and the estimated rates.

## Supplemental Figures

### Figure S1. Measured Point Spread Functions

R-Z (lateral-axial) images of 200nm beads obtained with SLAPMi’s line scan (top left) and raster scan modes (top center). The full-width half-maximum extents of the PSFs are:

Raster beam: 415 nm X, 1469 nm Z

Line beam: 432 nm X, 1617 nm Z

Without correction for the size of the bead. The corresponding diffraction limit for an ideal point source at NA 1.05 is 377nm X, 1423nm Z (Zipfel et al., 2003).

Line excitation results in slightly reduced axial sectioning because light is focused on only one lateral axis. As a result, the total intensity of the line focus decreases as 1/*z* away from the focal point, compared to 1/*z*^2^ for the point focus, where *z* is axial distance from the focal plane. Subtraction of the raster and line PSF highlights the axial side lobes in the line PSF (top right).

We incorporate a discrete convolution when using the image obtained with the raster beam to estimate measurements under the line beams, to account for these side lobes. The convolution kernel (example, bottom center) is estimated by deconvolving the line PSF with the raster PSF and constraining the solution (bottom left) to be positive and lie within the plane spacing of the reference image (0.75 µm). The raster PSF convolved with this kernel agrees well with the line PSF. Use of this model PSF improves agreement between expected and measured SLAPMi recordings (Fig. S6).

### Figure S2. SLAPMi excitation efficiency

Excitation efficiency (the number of fluorescence photons collected as a function of average excitation power) for SLAPMi and raster scanning, measured for fields of view with different degrees of sparsity. Recordings were performed in a fluorescent plastic block, with sparsity artificially imposed using the SLM. Photon rates were measured by summing photodetector current and dividing by the known mean current for a single photon. Lines are quadratic fits (forced 0-intercept) to the measured points. The absolute efficiency of raster scanning increases linearly with the density of labelled pixels. The absolute efficiency of SLAPMi is a peaked function of the label density, due to the imperfect extinction of excitation light by ‘OFF’ pixels of the SLM. This function reaches a maximum at a sparsity equal to the extinction coefficient of the SLM. We estimate that SLAPMi and raster scanning have equal efficiencies for fields of view with approximately 2% of pixels labelled. SLAPMi efficiency could be further improved by increasing the extinction coefficient of the SLM.

### Figure S3. Simulations of Activity Reconstruction

We evaluated performance of the solver by changing one parameter at a time using simulated data assuming the rest of the parameters are known to the solver. Segments used in the simulations were 10*10 pixel squares placed randomly (uniformly chosen) within a 500 * 500 pixel circle at the center of the field of view with 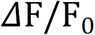 uniformly distributed in the 0.5-5 range. Where regularization was used, the algorithm enforced a maximum estimated 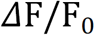 of 10 as the constraint. We performed 500 iterations of the dynamic Richardson Lucy algorithm.

The parameters used for different figures are listed in table below.

| Figure Name | # segments | $\Delta F/F_0$ | # random instances | # frames | Photons per frame per segment | $\tau$ | nshifts |
| --- | --- | --- | --- | --- | --- | --- | --- |
| S3-1_Corr_vs_ppp | 500 | Unif(0.5-5) | 5 | 350 | $10^{-1}$-$10^5$ | 100 | 0 |
| S3-2_Corr_vs_nseeds | 10-1000 | Unif(0.5-5) | 40 | 350 | 100 | 100 | 0 |
| S3-3_Corr_vs_tau | 500 | Unif(0.5-5) | 40 | 500 | 100 | 0-1000<br>(ground<br>-truth<br>=100) | 0 |
| S3-4_<br>Corr_vs_unsuspected | [500 750]<br>(500 gt) | Unif(0.5-5) | 40 | 500 | 100 | 100 | 0 |
| S3-5_Corr_vs_nshifts | 500 | Unif(0.5-5) | 40 | 350 | 100 | 100 | 1-10 |
| S3-6_Alternate_seg | 100-800 (see<br>fig) (20 each) | Unif(0.5-5) | 10 | 500 | 100 | 100 | 0 |
| S3-7_Corr_mtx | 500 | Onion method<br>(correlated F,<br>Unif(-0.6,0.6)) | 5 | 3000 | $10^{-1}$-$10^6$ | 100 | 0 |
| S3-8_ProjectionVs<br>Raster | Sparsity 1-20% | Unif(0.5-5) seed<br>activity | 20 | 1 | - | 100 | 0 |

**Evaluation metric:** We report the Pearson correlation between the estimated 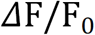 and its ground-truth counterpart over time for each segment, averaged over segments. Null distributions were obtained by randomly shuffling time points for each seed.

In the first simulation (Figure S3-1) we evaluated the effect of sample brightness on performance of the solver. This was done by fixing the *expected* Poisson rates per segment per frame to vary in the range 0.1-10^5^. As a comparison, typical values of this parameter in the recorded datasets in this work are around 100. Increasing the photon budget improves the reconstructions.

Figure S3-1: Effect of sample brightness on the solver.

In the second simulation (Figure S3-2), we changed the number of sources in the range 10-1000, in both the generation of the ground truth and solver. Increasing the number of sources decreases solver performance.

Figure S3-2: Effect of number of sources on the solver.

In the third simulation (Figure S3-3), we evaluated performance using different estimated decay time constants. The ground truth activity was generated using a time constant of 100 frames. We varied the time-constant in the range 0-1000. The solver performs best when the correct time constant is used. The solver is insensitive to underestimation of the time-constant, as this can be compensated with additional spikes (**w**) but does reduce the denoising benefit of the dynamics model. A time constant of 0 corresponds to normal Richardson-Lucy iterations. Substantial overestimation of the time constant degrades the performance of the solver.

Figure S3-3: a) the effect of erroneous decay time-constant on the solver. b) Null distributions for different time-constants.

In the fourth simulation (Figure S3-4), we evaluated the performance of the solver in the case where not all sources in the field of view were included in the segmentation. We added 0-50% additional unsegmented sources. This is meant to simulate fluorescence activity within the ‘ON’ region of the SLM but not detected in the reference image, a possibility in sensors with very low baseline fluorescence. Increasing unsegmented activity degrades the performance of the solver.

**Evaluation metric:** We report the Pearson correlation between the estimated

Figure S3-4: a) The effect of unsegmented activity on the reconstructions. b) Null distributions for different levels of unsegmented activity.

In the fifth simulation (Figure S3-5) we evaluated sensitivity of the solver to alignment errors. In order to do so, we shifted the reference image by 0-10 pixels after registration. Better alignment improves the reconstructions.

Figure S3-5: The effect of alignment (shifts in pixels) on the reconstructions.

In the sixth simulation (Figure S3-6) we evaluated performance of the solver when the reconstruction segmentations do not match the ground-truth activity segmentations. Alternate segmentations lead to a degraded performance which can be partially compensated for by using a finer segmentation than the ground truth.

Figure S3-6: The effect of alternate segmentations on the reconstructions.

We next quantified the undesired correlations introduced to the activities (Figure S3-7) as a result of the solver for the dynamic RL solver compared to the RL iterations. We generated 3000 frames of activity with random spiking patterns using vines and the extended onion method^41^, resulting in correlated and with correlations in the range −0.6 – 0.6. As can be noted, recovered activities using the dynamic RL iterations significantly improves.

Figure S3-7: recovered vs. ground-truth correlations. a) Dynamic RL iterations, b) RL iterations.

In the eighth simulation (Figure S3-8), we quantified the performance cost of mixed measurements, as compared to a hypothetical microscope that samples the volume at the same rate as SLAPMi, but can assign each photon to the correct source without the need for unmixing. This simulation provides intuition regarding the benefits of low degrees of multiplexing. We show performance of the two methods for sparsity levels in the range 1-20 % and sample brightnesses of 0.1-100 photons per source per frame.

Figure S3-8: performance of SLAPMi vs raster scanning for different sparsity levels and photon budgets

### Figure S4. Automated Segmentation

The goal of our automated segmentation procedure is to group pixels such that each segment within the SLM ‘ON’ area belongs to a single neuronal compartment (such as a spine), all segments are of a minimum brightness, and the total number of compartments within the SLM ‘ON’ area is less than 1000. These constraints can only be approximately satisfied. To do this, the reference image (left, gamma=0.5) is classified into 4 classes (center) using a hand trained classifier in Ilastik’s autocontext mode. The classes are meant to correspond to dark (background) voxels, spine heads, voxels with fluorescent label, and dendritic shafts. The voxel classes and image intensity are passed to a skeletonization-based algorithm that groups non-dark pixels into segments. The automated segmentation output (right) occasionally produces flaws such as fused or split spines, which we have not yet quantified. We have written an interface for manual curation of these errors, which was used for most datasets. See the published code for more details.

### Figure S5. *In Vivo* Fast Motion Correction

Recorded SLAPMi data before (left) and after (right) cross-correlation based fast motion correction. This is a magnified view of just a portion of one line scan. In this recording measurements were spaced by 125 nm. Low-amplitude motion is ubiquitous in mammalian brain imaging, due to blood flow and animal movements. SLAPMi imaging at 1 kHz is considerably faster than these sources of movement, allowing compensation by rigid registration over the fields of view imaged here without ‘rolling shutter’ artifacts.

### Figure S6. Global Position Alignment

SLAPMi measurements (average of 100 frames) and Expected SLAPMi measurements before (top) and after (bottom) global position alignment. The Expected measurement is the product of the measurement matrix (**P**) with the rigidly-aligned reference image. Alignment is performed minimizing a distance metric between measured and expected signals over a range of 3-dimensional offsets using multiscale grid search.

### Figure S7. Pixelwise analysis of jRGECO and yGluSnFR tuning in cortical dendrites

**a)** Correlation between tuning curves measured yGluSnFR and jRGECO channels as a function of jRGECO response amplitude, for pixels showing significant stimulus responses in the jRGECO channel (p<0.01, t-test) that also exceeded a threshold brightness in the yGluSnFR channel. Highly responsive pixels in the jRGECO channel, corresponding to responsive spines, show similar tuning curves in both channels.

### Figure S8. Conservation of global modes across stimuli

(top) Spatial correlations between principal components obtained separately for trials of each stimulus type, in an example session. The largest components in each data subset correspond across subsets. (bottom) correlation of corresponding temporal weights to estimated motion magnitude.

### Figure S9. SiPM detector performance

(top) Voltage trace recorded from detector during SLAPMi acquisition. Only a brief period of time is shown here. Digitization (250 MHz) is synchronized to laser emission (5MHz) via a phase-locked loop. On each laser pulse, an integer number of photons arrives at the detector, resulting in distinct response amplitudes. (bottom) Pulse height spectrum of the detectors. The histogram of detector response amplitudes (normalized to the single-photon amplitude) shows distinct peaks that allow the number of simultaneous photons to be accurately estimated from the photocurrent. The pulse height spectrum for a photomultiplier tube, by comparison, does not show distinguishable peaks for more than 2 photoelectrons, and hybrid photodetectors represent a much smaller improvement in this respect (see “Hybrid photodetectors combine PMT and semiconductor diode technologies.” Motohiro Suyama, Hamamatsu Photonics K.K., and Maridel Lares, Hamamatsu Corporation; 2008).

